# Subcortical control of reaching in humans

**DOI:** 10.1101/2025.11.26.690644

**Authors:** Samuele Contemori, Gerald E. Loeb, Brian D. Corneil, Guy Wallis, Timothy J. Carroll

## Abstract

Accurate visually guided reaching requires transformation of target-related photoreceptor responses into precisely coordinated activation of trunk and arm muscles. The cerebral cortex is widely believed to compute the requisite kinematic and musculoskeletal dynamics strategies in humans ^1–3^, even though vertebrates lacking a cerebral cortex achieve sophisticated visuomotor control ^4–6^, and brainstem circuits executing coordinated eye and head gaze shifts perform analogous sensorimotor computations in non-human primates ^7^. Here we used a visuomotor reaching task that yields extremely rapid, “express”, target-directed muscle activations ^8–10^ to test whether a putative subcortical sensorimotor network can compute musculoskeletal dynamics to initiate reaching in humans. We found coordinated express visuomotor responses (EVRs) in task-relevant shoulder, elbow, and bi-articular muscles that reflected both starting posture and target direction in similar patterns to longer latency, presumably cortically mediated, muscle responses. When the task goal was to reach away from the stimulus (i.e. an “anti-reach”; ^11^) the EVR involved coordinated muscle activation to initiate the hand toward the stimulus location, opposite to the subsequent goal-directed response. The results suggest a unified theory of visuomotor control for reaching and gaze shifts, in which subcortical systems compute musculoskeletal dynamics based on sensory target information and cortically derived context. The results imply that the transformation from motor goals in extrapersonal space into musculoskeletal dynamics can be performed by neural circuitry in humans that does not involve the sensorimotor cortex.

## MAIN BODY

The capacity for fast and accurate visually-guided movements emerged at a primitive stage of animal evolution. Consider the impressive prey capture behaviours of early diverging vertebrates such as frogs and archerfish, which lack a cerebral cortex and yet can catch a fly with apparent ease. The brainstem and spinal circuits that drive such behaviour have been phylogenetically conserved in primates ^5,6^, where they are known to account for both voluntary and express orienting behaviours of the eyes and head ^7,12^. Despite long-standing evidence for multiple descending systems that converge upon spinal interneuron circuits and are capable of considerable sensorimotor integration ^13–15^, studies of visually guided human limb movement have usually considered only the role of the corticospinal system. This likely stems from the difficulty in experimentally accessing brainstem structures in humans with sufficient spatiotemporal resolution to provide evidence about their functional roles.

Striking, but generally overlooked, parallels exist between the control networks for gaze and limb movements. The frontal and supplemental cortical eye fields are adjacent to the primary and secondary motor cortices and both cortical systems are necessary for planning kinematic trajectories for voluntary behaviours such as tracking objects for the eye fields and reaching to objects for the motor cortex ^16–18^. Both eye and limb related cortical areas have strong projections to the superior colliculus and brainstem premotor nuclei ^19–21^, which also receive multimodal sensory information about potential targets directly via subcortical pathways, thereby enabling express behaviours ^22,23^. The muscle activations required for dynamic control of gaze require coordination between eye and head movement and are computed in those subcortical structures rather than relying on direct corticomotoneuronal projections ^7^.

Here we examine the capacity of human subcortical control systems to compute the dynamic patterns of muscle activations needed to accurately drive the limb toward visual targets. We do this via a behavioural manipulation that yields extremely rapid target-directed muscle activations that are unlikely to be generated cortically ^9,10,12,24–28^. We term these short latency muscle activations “express visuomotor responses” (EVRs), to draw a parallel with express saccades. EVRs appear as brief bursts of electromyographic (EMG) activity at a narrow set of onset times locked to target appearance (∼70-120ms), irrespective of the latency of subsequent EMG bursts (i.e. long latency responses; LLRs) or mechanical reaction times (RTs), suggesting that two distinct control signals reach the muscle. Larger EVRs are associated with faster movement initiation ^9,24,25^, but EVRs lack strategic flexibility in that they are always directed to the physical location of the stimulus, even if the instruction is to reach in the opposite direction (i.e. for anti-reaches ^25^). Thus, express visuomotor responses closely parallel express saccades, which are exclusively stimulus-directed rather than goal-directed in the well-studied anti-saccade task ^29,30^. Here we test whether EVRs exhibit the same multi-muscle, posture-dependent coordination patterns as the longer latency, presumably cortically mediated, LLRs. If so, it would support the hypothesis that similar subcortical systems are used to compute the dynamics of the required muscle activations for both voluntary and express gaze and reach behaviours.

### Distribution of Express and Voluntary Muscle Activity

We first asked whether the control system for express visuomotor behaviour can account for limb mechanics to produce muscle activity that drives the hand toward visual targets under a diverse range of conditions. Our hypothesis was that EVRs reflect a fully coordinated control signal that appropriately drives all the major shoulder and elbow muscles required for target-directed reaches in different directions in extrapersonal space and from different starting postures. We recorded the activity of eight proximal and distal arm muscles while participants performed reaches to eight radial targets, using a moving target protocol that is known to promote EVRs (^24,28^; Figure 1b). The same eight reaches were performed in two different shoulder postures (abducted and adducted; Figure 1a) that were expected to require small but systematic differences in muscle activation patterns for the same hand trajectories ^18^.

**Figure 1:**
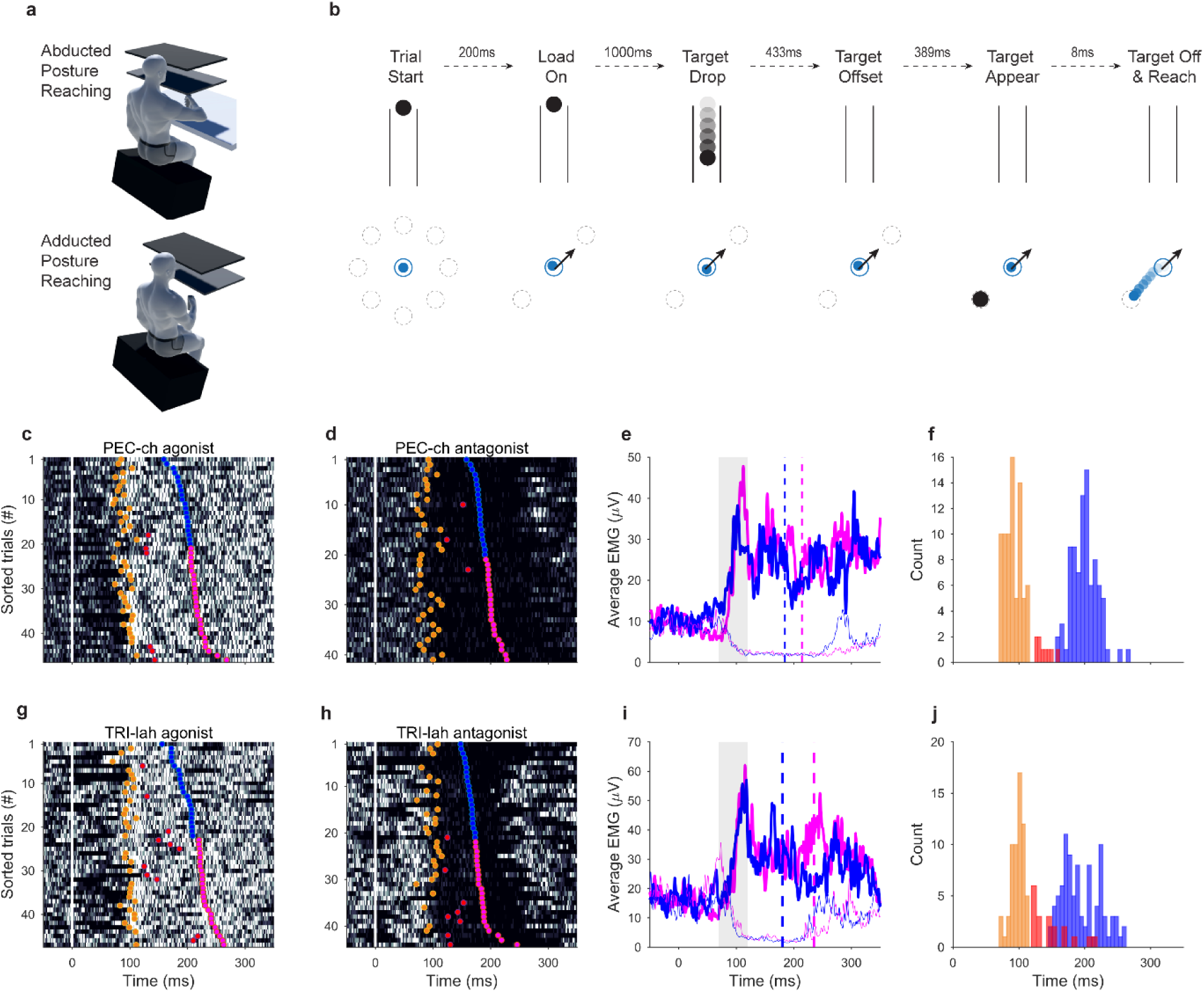
**a,** Schematic of the abducted and adducted shoulder postures. In the abducted-shoulder posture, the upper limb was supported on a custom-built air sled positioned under the right elbow to minimize arm-on-table sliding friction (not shown here). **b**, Schematic of the moving target paradigm. The reaching hand position was virtually represented via a cursor (blue dot) displayed on the monitor and projected into the (horizontal) plane of hand motion via a mirror. To start the trial, participants had to keep the cursor at the ‘home’ position resisting the direction-dependent robot-force (black arrow), which preloaded the arm muscles and approximately cued the two possible target locations on each trial. In this example, the robot force has a 45° orientation so the target could appear only at 45° (toward the load) or −135° (away from the load). **c,d** Raster plots of surface EMG activity of the PECch muscle of an exemplar subject who performed target-directed reaches toward a −45° target (**c**), or a 135° target (**d**) in the adducted shoulder posture. The robot pre-loaded the upper-limb with a force oriented to the 135° direction. Greater EMG amplitude is shown by lighter shading in each raster, the white vertical line at 0ms indicates the target onset time, the orange scatters represent the initiation times of express muscle responses, red scatters show EMG onsets outside the express window, blue scatters indicate hand movement onsets with RT<median, and the magenta scatters shows hand onsets with RT>median. **e**, shows the average EMG activity when the muscle was agonist (thick traces) or antagonist (thin traces) for the required target-directed reach for the slow (magenta) and fast (blue) trial subsets (+/− median RT). The grey patch defines the express response time window (i.e. 70-120ms after the stimulus onset) and the dashed vertical lines represent the average RTs for each subset. **f**, histogram of muscle activation onsets (orange = express, red = non-express) and hand RTs (blue). **g-j**, same as c-f but for the TRI-lah during reaches toward a 0° target or a 180° target while the robot loaded the upper-limb in the 180° direction.

Figure 1 shows EMG traces from muscles acting at the shoulder (Pectoralis Major clavicular head; PEC-ch, Figure 1c-f) and at the elbow (Triceps Brachii lateral head; TRI-lh, Figures 1g-j) during target-directed reaches for an exemplar participant. Express visuomotor responses appear as a vertical band of muscle activations when the muscle was agonist for the movement (figure 1c,g), or inhibitions when the muscle was antagonist (figure 1d,h). The onset time of express responses (brown scatters in figure 1c,d,g,h) was consistently at ∼90-100ms from the target presentation, irrespective of the RT defined by the initiation of hand motion (see the blue vs magenta traces for RTs greater or less than the median RT in figure 1e,i), which is the signature of two distinct control signals reaching the muscle. The mean EMG amplitude in the express response window was substantial compared to subsequent “voluntary” muscle activity, for trial subsets with both early and late hand movement onset times. Across participants and muscles, the average prevalence of trials in which target-directed muscle activity was detected in the EVR window was 51±5%, and the mean onset latency ranged from 88ms to 93ms for proximal to distal muscles (see supplementary table 1). This indicates that the express visuomotor control network can rapidly and substantially recruit both proximal and distal arm muscles at short and consistent latency, as would be expected for a short neural pathway linking visual inputs to target-directed reaches.

We found similar modulation of normalised (see methods) EVR and LLR muscle activity across targets for each of the eight recorded muscles (six are shown in Figure 2), suggesting that similar patterns of agonist-antagonist muscle coordination were recruited during the two temporally-distinct epochs. Because EVRs are facilitated by muscle preloading ^9,25,31^, we were primarily interested in trials in which the target location was opposite to the direction of background force applied by the robot (figure 2). However, the target could randomly appear either toward or opposite to the robot force direction to prevent target predictability, so we also measured EVRs in the trials when muscles were not pre-loaded (supplementary figure 1). Notably, muscle activity patterns were similar between EVRs and LLRs irrespective of whether the muscle was loaded prior to target appearance, thus excluding the possibility that the direction of preload bias drove the target-dependent muscle activity. Note also that EVRs in a single muscle are tuned to stimulus direction when the upcoming target is unpredictable among eight radial targets ^9,32,33^.

**Figure 2.**
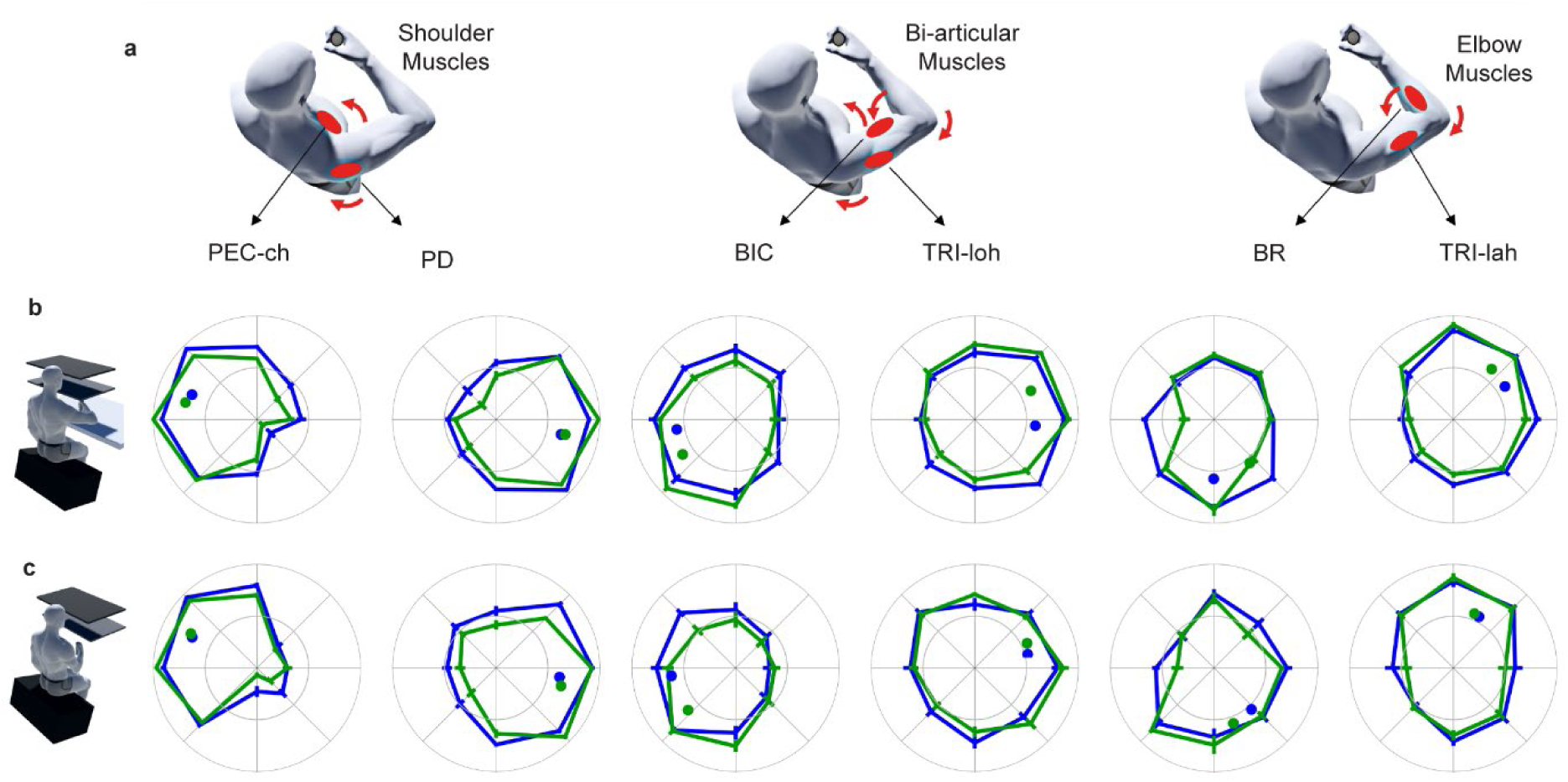
**a**, Schematic of the location and function of the six muscles used to illustrate target- and posture-specific muscle activation patterns for express and long latency muscle responses. **b**, Express (blue) and long-latency (green) muscle activity for each target direction in the abducted posture. Each radial plot shows average (+/− SE) target-dependent tuning curves across the twenty participants. Express and long-latency activation values are normalised to the maximal EVR and LLR amplitudes respectively across target directions. The outer ring indicates the normalised value (1) and the inner ring represents the reference background activity (0) for each direction. Thus, excitations with respect to baseline lie between the two rings, and inhibitions from baseline fall within the inner ring. Blue (EVR) and green (LLR) circles show the preferred direction of each tuning curve. **c**, Same as **b**, except for the adducted posture. PEC-ch: Pectoralis Major clavicular head, PD: Posterior Deltoid, BB: Biceps Brachii, TRI-loh: Triceps Brachii long head, BR: Brachioradialis, TRI-lah: Triceps Brachii lateral head.

We next asked whether the pattern of multi-muscle coordination for express responses was functionally modulated by the starting posture of the limb (see figure 2b versus 2c). As expected ^18^, we found that some, but not all, muscles showed changes in directional tuning with posture shifts for muscle activity in the long latency response window. Specifically, the preferred direction of the normalised LLR amplitude (i.e. from the vector sum from the 8 directions) was significantly different between the adducted and abducted postures for five of the eight muscles sampled (Pectoralis clavicular and sternal heads, Posterior Deltoid, Brachioradialis, Triceps lateral head; Watson-Williams test ^34^; see supplementary table 2). There was a significant corresponding change in EVR preferred direction for three of these five muscles (Pectoralis clavicular head, Brachioradialis, Triceps lateral head), and the trend in preferred direction changes was the same for EVRs and LLRs in all eight muscles (see figure S1).

We next ran a principal components analysis (PCA) to formally test whether the patterns of multi-muscle coordination observed in the EVR were comparable to those evident in the LLR for the same limb posture. We reasoned that if the EVR embodies a fully coordinated control signal that appropriately drives all the major shoulder and elbow muscles to targets, then it should be possible to reconstruct the muscle activities observed in the LLR using a low-dimensional subspace that was computed from the EVR. Crucially, if the EVR is appropriately modulated to compute the muscle coordination patterns needed for specific starting limb postures, then reconstructions of the LLR activity from the EVR in the same posture should be more accurate than reconstructions of EVRs or LLRs from the alternative posture. To test this, we ran a PCA on EVR muscle activities recorded in the abducted posture and compared how well this low-dimensional subspace could account for muscle coordination observed in the LLR in abduction and both the LLR and EVR in adduction. The first four principal components (PCs) extracted from the abduction EVR accounted for over 95% of the variance (VAF) in the sampled EMG activity on average across the group (See figure 3a). A 2-way rmANOVA revealed that the VAF was significantly modulated by the posture condition (abducted vs adducted; F_1,19_=41, *p* <0.001) and the interaction of posture with response type (F_1,19_=40.6, *p* <0.001), whereas there was no statistically significant effect for response type (express vs long-latency; F_1,19_=2.2, *p* = 0.15). The post-hoc analysis showed that the low-dimensional subspace extracted from the EVR interval was sufficient to explain ∼90% of the variance of long-latency muscle activity in the same posture (Figure 3a), yet significantly less capable of reconstructing either the express or long-latency muscle activity in the adducted shoulder posture. To exclude the possibility that these results simply arise from short RT trials where the EVR window is contaminated by EMG from a long-latency control signal, we confirmed that similar results were found for data subsets with more restrictive minimum RT cut-offs (i.e. RT > 170 and RT > 200ms, see fig S2). To exclude the possibility that background load related EMG could explain the observed EVR coordination patterns, we confirmed that similar results occurred for muscle responses recorded in unloaded muscle conditions (see fig S3). Overall, the results confirm that posture-dependent muscle coordination patterns were similar for express and long latency responses.

**Figure 3.**
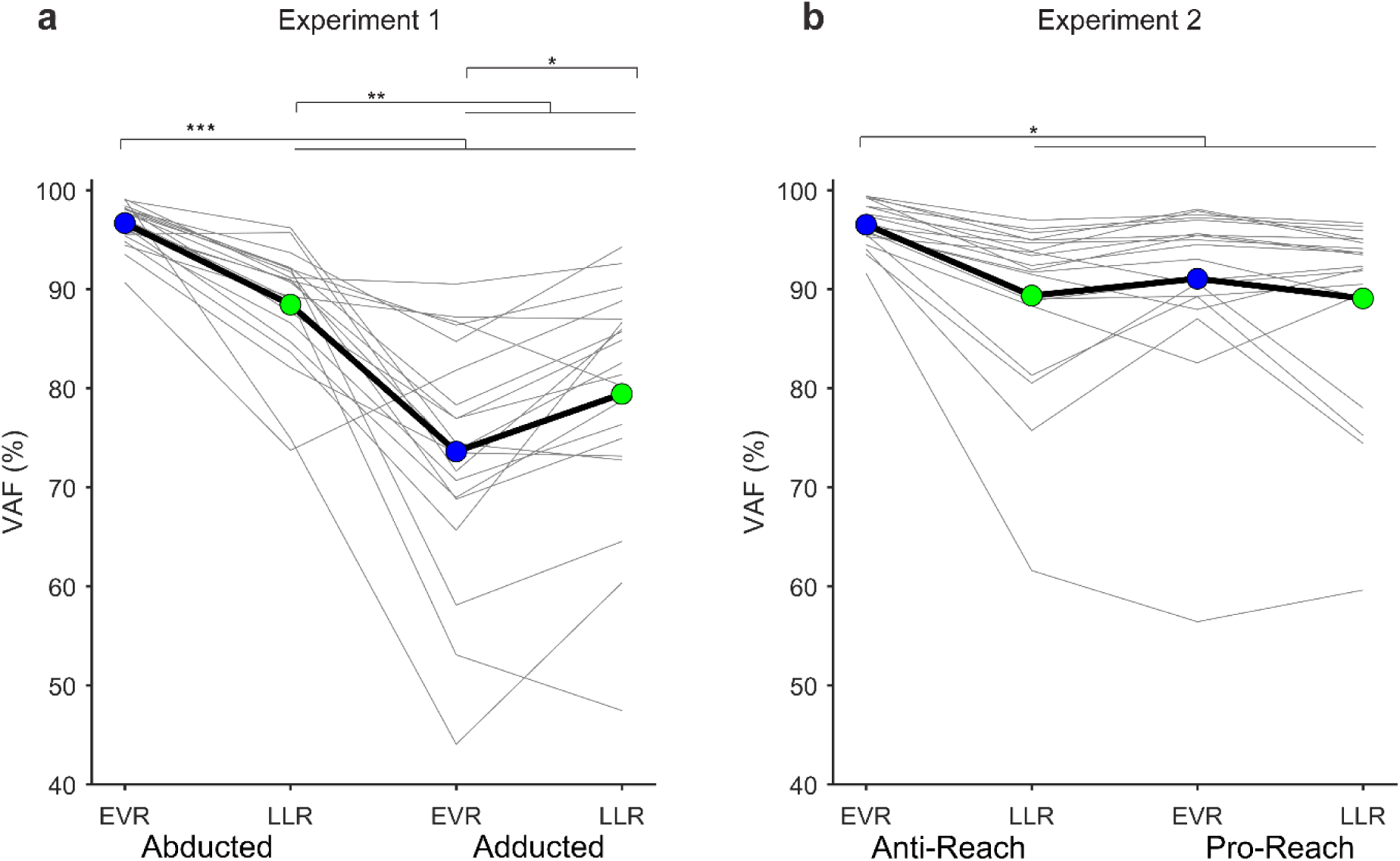
**a,** Variance accounted for (VAF) by the top 4 principal components extracted from the EVRs across stimulus directions in the abducted shoulder posture, for LLRs in both postures and EVRs in the adducted posture in experiment 1. **b,** VAF by the top 4 principal components extracted from EVRs in the anti-reach condition, for LLRs and EVRs at the same stimulus location in the pro-reach condition, and LLRs in the opposite direction for the anti-reach condition in experiment 2. EVR means are blue, LLR means are green. The brackets and asterisks indicate the statistically significant (* *p*<0.05, ** *p*<0.01, *** *p*<0.001,) post-hoc pairwise contrasts.

The most rigorous test of the functional capacity of the control system for express behaviour is when the goal of the reach is different from the target stimulus location. In this way, the tuning of both the EVR and the LLR can be tested against either the stimulus or the goal location. If the multi-muscle coordination of the EVR is accurately tuned towards visual stimuli in multiple directions irrespective of the goal location, it would provide strong evidence that a low-level system that is incapable of abstract stimulus-response mappings can nonetheless compute complex musculoskeletal dynamics. We therefore ran an experiment in which we asked participants to reach either toward (pro-reach) or in the opposite direction from (anti-reach) a set of eight stimulus locations. The pro-reach and anti-reach trial subsets were randomly intermixed and designated on each trial by cursor colour changes ∼1s before stimulus presentation. Unsurprisingly, the higher complexity of the anti-reach task led to longer RTs relative to pro-reach conditions (Supplementary figure 2; group-level RT results: pro-reach 236±23ms, anti-reach 292±23ms, paired t-test *p*<0.001). Crucially, however, the onset-time of express responses was independent of the task requirements (∼85ms for PECch, ∼100ms for TRIlah, in the exemplar participant in Figure 4). Whereas both express and long-latency muscle responses accurately encoded stimulus locations in all directions under pro-reach conditions, the anti-reach trials led to opposite tuning directions for express and long-latency responses (Figure 4). The results show that the sensorimotor control system that generates EVRs strictly converts hand-to-stimulus discrepancies into stimulus-directed muscle activation at short-latency, irrespective of the task rules. This provides further evidence that express and long-latency responses reflect distinct motor control signals; an early signal that invariably encodes the stimulus location (EVR) and a later signal that is flexible to contextual task rules (LLR). We presume that the LLR is cortically mediated because humans with frontal lobe lesions including the cortical eye fields are incapable of correctly performing anti-saccades ^35^. In non-human primates, the supplementary eye fields appear crucial for the transformation from a stimulus location into a spatially incongruent goal location for anti-saccades ^36^, and the dorsal pre-motor cortex likely contributes an analogous function for anti-reaches ^37,38^.

**Figure 4.**
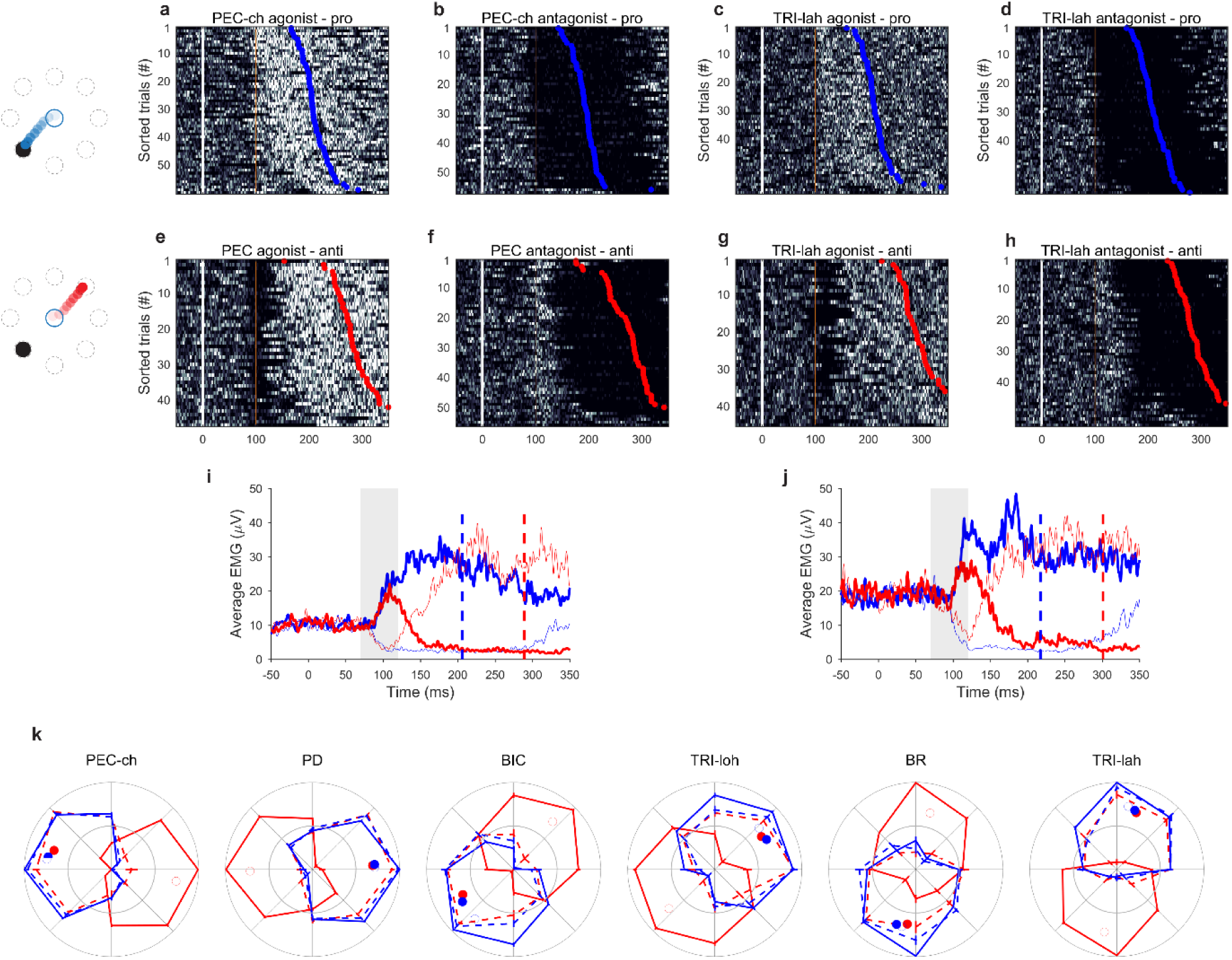
Raster plots of surface EMG from the PEC-ch muscle of an exemplar subject who performed goal-directed reaches toward a −45° reach target (**a,e**), or a 135° reach target (**b,f**) in pro-reach (**a,b**) and anti-reach (**e,f**) conditions (same format as figure 1). Thus, the visual stimulus locations were opposite for the pro- and anti-reach conditions in these plots. The orange vertical line at 100ms gives a visual reference to illustrate express response onset-time. **i** average EMG for pro-reach (blue) and anti-reach (red) trials for visual stimulus locations when the PEC-ch was agonist (thick traces) and antagonist (thin traces). **c,d,g,h,j** same for the TRIlah muscle during reaches toward a 0° reach target or a 180° reach target while the robot loaded the upper-limb in the 180° direction. **k**, Express (solid lines) and long-latency (dotted lines) target-dependent muscle activity in six shoulder and elbow muscles (same format as Figure 2). Red lines represent anti-reach trials and blue lines show pro-reach trials.

We next asked if the express muscle activity that appeared oriented to the physical stimulus location in the anti-reach conditions represents a functionally coordinated response across all muscles (Figures 3b, 4k). We tested if the express and long-latency muscle coordination of pro-reaches could be reconstructed using the PCs extracted from the anti-reach EVR for the same visual targets. We also tested if the anti-reach LLR muscle coordination could be reconstructed from the anti-reach EVR PCs for targets in the opposite direction. Although the VAF was significantly modulated by task condition (pro-reach vs anti-reach; F_1,19_=9.8, *p* =0.006) and response (express vs long-latency; F_1,19_=16.8, *p* <0.001), and the task*response interaction was marginal (F_1,19_=9.4, *p* =0.006), in all cases the VAF from the EVR-ANTI trials explained over 90% of the variance for the other trial types. The small drop in VAF seen across the different trial conditions in Experiment 2 (Figure 3b) is comparable to the drop observed between EVRs and LLRs within the same posture in Experiment 1, and far less than that observed across postures (Figure 3a). The results show that 1) similar patterns of muscle activation and inhibition were recruited for each visual stimulus location during the EVR response phase irrespective of the ultimate reach goal, and 2) these muscle activity patterns were similar to those associated with the subsequent LLR muscle activation that initiated the limb towards the corresponding goal location (i.e. when reach directions are inverted for the LLR in the anti-reach condition).

## DISCUSSION

### Neural Substrates of the Express Visuomotor Response

The data show that whatever neural circuits underlie EVRs, they are capable of producing coordinated muscle activity in shoulder, elbow and bi-articular muscles to accurately reach towards visual stimuli: irrespective of starting posture, whether or not the muscles are pre-loaded, or whether the task goal is to reach towards or away from the stimulus location. We propose that these computations are performed by the tecto-reticulo-spinal system, in a direct parallel with the role of subcortical control components of the oculomotor control system for saccadic gaze shifts ^7,12,39^. Here we briefly summarize three convergent lines of evidence for this hypothesis (see ^8–10,24–28,31,40–42^ for detailed arguments).

First, the short latency of EVRs (∼90ms on average in this study) implies a short transmission path that precludes extensive processing within sensorimotor cortex. The minimal latency for target-related activity within motor cortex is difficult to estimate precisely in humans but is likely to be ∼50ms in non-human primates ^43–45^. When compared to express muscle activation in rhesus monkeys ranging from 48-91ms (mean ∼65ms) on deltoid muscles during reaching ^42^, or 56-97ms (mean ∼73ms) in the neck during gaze shifts ^41^, it is clear that opportunity for motor cortical processing of visual inputs prior to EVR onset is minimal. Note that a ∼30ms loop time, from receipt of sensory input in M1 to the onset of muscle activity, is required for cortical contributions to the long latency stretch reflex ^46^, and that closer to 100ms from stimulus onset is required for reliable decoding of visual target direction within M1 (125 ± 28, ^47,48^), or occupation of visual target related dimensions within the execution subspace of neural state (71-114ms, ^49^).

Second, there are striking similarities between the properties of eye and neck components of express gaze shifts and limb EVRs, in terms of timing, preferred stimulus features, and limitations in flexibility to task context ^8,10,24,25,27,28,30,40–42,50–52^. There is definitive evidence in non-human primates that express saccades are driven by the earliest arrival of visual input to the superior colliculus and subsequent recruitment of the brainstem oculomotor system ^39,51,53,54^, prior to contributions from frontal or supplementary eye fields. Similarly, visual responses in the superior colliculus precede neck muscle EVRs by a latency that resembles the efferent lag from the superior colliculus rather than frontal eye fields ^55–57^. It also appears that a retino-tecto-reticular pathway remains functionally viable for saccade modulation in humans following hemidecortication ^58^.

Third, the brainstem and spinal cord have the computational capacity to convert motor goals into musculoskeletal dynamics. At the spinal cord alone, neuromechanical simulations show that muscle afferents and associated spinal interneuronal circuitry are sufficient to transform discrete “set” and “go” input signals into coordinated time-varying muscle activations for target-specific multi-joint reaching ^59^. Physiologically, the tecto-reticulo-spinal control system is the dominant system for visually-guided behaviour in early diverging vertebrates that lack a cerebral cortex ^4,5,60^, but it has also long been known that subcortical pathways are sufficient to generate goal-directed behaviours in mammals. Decorticate dogs and cats execute goal-directed movements of their head and limbs in response to stimuli of various sensory modalities ^61,62^. More recent work using powerful genetic and viral tracing techniques has identified specific brainstem circuits sufficient for complex forelimb movements in mice ^63,64^, including those involving the superior colliculus ^65^. Crucially, the brainstem is capable of coordinating visually-guided limb function in primates; macaque monkeys can run, jump and climb within days of complete bilateral pyramidotomy ^66,67^. In humans with strokes and macaques with pyramidal tract lesions, recovery of motor function is associated with changes in the reticulospinal subsystem ^68–70^. These have been identified using transcranial magnetic stimulation ^71^ and magnetic resonance imaging ^72^ based on ipsilateral reticulospinal projections (as opposed to the contralateral projections of motor cortex). Finally, similar to the EVRs presented here, startling acoustic stimuli accompanying a reaching cue shorten RTs at latencies that preclude a cortical pathway ^73^. This StartReact effect increases following stroke ^70^ and has been used to argue for an involvement of the reticulospinal subsystem in primate reaching ^74^. Crucially, combining startling acoustic stimuli with visual targets for reaching increases the magnitude of EVRs without affecting their latency ^26^, suggesting a spatiotemporal summation of visual and auditory signals within the brainstem (see also ^75^).

### Accounting for Cortical Activity in a Distributed Movement Control System

If the musculoskeletal dynamics of at least some visually guided behaviours are computed subcortically, then what is the role of cortex in those behaviours? Neural activity in motor cortex appears to anticipate musculoskeletal dynamics ^1,18,76^, and such activity is widely interpreted to define the state of a dynamical system that generates movement commands (e.g.^1,14,76–78^). However, the current data show that the transformation from motor goals in extrapersonal space into musculoskeletal dynamics can occur at latencies that likely preclude a typical evolution of motor cortical neural state through preparatory and execution sub-spaces ^49^. This fits our proposal that EVRs are generated subcortically, but it is an open question whether the presumably cortically driven long-latency responses exert their effects 1) in parallel to the putatively subcortical control system that produces EVRs (e.g. via the corticospinal tract), or 2) through the control system responsible for EVRs (i.e. via a cortico-reticulo-spinal pathway), as is the case for gaze saccades.

The functional sophistication of the EVR control system emphasises that the motor cortex is part of a hierarchical ^13,15,79^ (or heterarchical ^14^) dynamical system that includes subcortical networks capable of complex sensorimotor transformation. Crucially, this system is bi-directional, in that each module both sends and receives information to and from interconnected areas ^79^. Thus, one role of preparatory cortical activity might be to alter the states of subcortical modules in anticipation of subsequent sensory or voluntary movement initiation inputs (e.g. an alternative interpretation of ^80^), while one cause of execution-related cortical activity might be efference copy from subcortical command centres in anticipation of anticipated afferent feedback ^81^. Indeed, rapid externally imposed movement that generates spinal stretch reflexes results in phasic excitation of neurons in both M1 and reticular formation that is modulated by the spinal reflex magnitude independently from the stimulus magnitude ^82^. Such efference copy information of the spinal reflex output would be invaluable to inform subsequent, cortically mediated corrective responses. Crucially, for long latency responses from motor cortex to be accurate, whether to imposed limb perturbations or the visual stimuli considered here, they must take into consideration the anticipated and actual direct contributions of subcortical centres such as the spinal cord and reticular formation. Such effects occur within the oculomotor system, where efference copy outputs from superior colliculus are transmitted to the cortical eye fields to allow cortical compensation for initial saccade endpoints during double step saccade tasks ^83^. Parsimony would suggest similar corollary inputs to motor cortex should be expected from the circuits generating the EVRs observed here.

Previous studies of EVRs in humans documented how contextual information about reaching tasks can modulate the behaviour of the putative tecto-reticulo-spinal system, presumably via cognitive cortical output projections ^10,24,31^. Cortical areas such as the frontal eye fields and pre-frontal cortex provide similar cognitive modulation of the tectal pathways responsible for gaze behaviours ^29,84^. Memory of prior performances and the efforts and rewards associated with them would motivate strategic decisions about participating in new tasks and seeking improved performance. The current results emphasise that the human cerebral cortex implements such strategic decisions by altering the states of relevant subcortical control systems, rather than just the final common pathway of the motoneurons.

## MATERIALS AND METHODS

### Participants

Twenty-five healthy adults were recruited for this study, which involved two experiments. Fifteen people completed both experiments, and each experiment had a total sample of 20 (Experiment 1: 11 males, 9 females; mean age: 27.4±5.7 years; Experiment 2: 10 males, 10 females; mean age: 27.3±7.6 years). All participants were right-handed, had normal or corrected-to-normal vision, and reported no current neurological or musculoskeletal disorders. They provided informed consent and were free to withdraw from the experiment at any time. All procedures were approved by the University of Queensland Medical Research Ethics Committee (Brisbane, Australia) and conformed to the Declaration of Helsinki.

### Experimental set-up and task design

The participants performed target-directed reaches using a two-dimensional planar robotic arm (vBOT) ^85^. The task paradigm was created in Microsoft Visual C++ (Version 14.0, Microsoft Visual Studio 2005) using the Graphic toolbox, and projected to the participants via a mirror that reflected an LCD computer monitor (120Hz refresh rate) and occluded direct vision of the arm (Figure 1a). In the vBOT display, the reaching-hand position was virtually represented by a cursor (1 cm diameter) whose apparent position coincided with the actual hand position in the plane of the robot motion. For the first experiment, the cursor was coloured blue, and the participants performed target-directed reaches from abducted and adducted shoulder postures (Figure 1a). In the abducted posture condition, the reaching arm was supported on a custom-built air sled positioned under the elbow to minimize arm-on-table sliding friction. For the second experiment, the participants were asked to perform either pro-target or anti-target reaches from the abducted shoulder-posture. The cursor color-coding cued the contextual task rule (green → pro-reach; red → anti-reach) from the trial start (Figure 4), approximately 1.5s before target onset.

For all experiments, target stimuli were a filled black circle of 3 cm diameter presented against a light grey background that could appear at one of eight possible radial locations 7 cm from a central ‘home’ position (Figure 1b). The target was presented via a *moving target* paradigm (Figure 1C) ^24,28^. In this paradigm, the target was initially displayed at the top of the monitor concurrently with a blue ring of 2 cm in diameter at the centre of the monitor, which indicated the starting “home” position of the hand. To start the trial, the subjects had to bring the cursor at the home position and to keep it there for ∼200ms as the robot gradually applied a load to the reaching arm (see below for details). Note that an additional period of 500ms was provided in the second experiment to ensure unambiguous extrapolation of the cursor color-coded task instructions. The participants had to counteract the robot force by keeping the cursor as stable as possible at the home position for ∼1s. The target then dropped at 30cm/s for 13cm along a vertical track (433ms), disappeared for 389 ms, and briefly reappeared (for 1 screen frame - ∼8 ms) at the stimulus location for reaching. This allowed us to generate a transient and temporally predictable stimulus that has proven effective to facilitate EVRs ^24^. The time at which the stimulus physically appeared (time 0 in all results) was recorded with a photodiode that detected a secondary light that appeared simultaneously with the target at the bottom-left corner of the monitor. The photodiode fully occluded the secondary light, thus making it invisible for the participants.

The pre-target loading force exerted by the robot had constant magnitude (5N maximum) and was modulated to account for direction-dependent force capability ^86^. Note that the robot-force direction cued the possible target locations, such that the final target could appear with equal probability either congruently with or opposite to the robot-force direction (Figure 1b). The preload, therefore, enhanced the background activity of the agonist or antagonist muscles required to reach the target. Importantly, both muscle preload and temporal predictability of the target are relevant features that facilitate the generation of detectable express visuomotor responses ^25,40^. EVRs for individual muscles are also tuned appropriately when targets are randomly drawn from larger sets of target locations ^9,32,33^.

The participants were instructed to start reaching to the target as soon as they saw it. “Too fast”, or “Too slow”, errors were shown if participants left the home position before target presentation or more than 500ms after target presentation, and the trial was repeated at the end of the trial block. We also asked the participants to end their movements at the target location because we recently found that express behaviour is impaired by deliberately overshooting or undershooting the target ^10^.

For the first experiment, participants completed two sessions, one for each posture. Each session involved 800 reaches comprised of 10 trial blocks (80 reaches/block). Each trial block contained 10 reaches to each of the eight target-locations, half of which had the robot pre-load aligned with target direction, and half with the pre-load opposite the target direction. All trials were randomly intermingled within blocks. For the second experiment, subjects completed 20 identical trial blocks in the adducted posture across two separate sessions. Each trial block contained six repetitions to each of the eight target-locations (half with pre-load aligned, half with pre-load opposite) for both pro- and anti-reach conditions, with all trials randomly intermingled (i.e. 96 reaches/block or 960 reaches each for the pro- and anti-reach conditions across the two sessions).

### Data recording and analysis

#### Kinematic data recording and reaction time detection

Reaching kinematic data were recorded via the vBOT optical encoders at a sampling rate of 1 KHz. To identify the movement reaction time (RT), we first defined the first hand-velocity peak exceeding the average value across the 100 ms prior to the target presentation by more than five standard deviations. We then fitted a line to the hand-velocity data enclosed between 25% and 75% of the peak velocity and indexed the RT as the time point when this line crossed the baseline (zero) velocity value ^10^. We also defined the initial reach direction by taking the slope of a line connecting the hand position coordinates at the RT and at 75% of the peak velocity. For the first experiment, we discarded trials in which the RT was <140 ms (3.5±3.9%) to minimise the possibility of temporal overlap between EVR and LLR signals, or the initial reach direction diverged by ±30° from the reach goal location (3.2±2.3%). For the second experiment, the trials were respectively classified as correct pro-reach, or anti-reach, if the hand-velocity vector at the peak velocity was directed toward, or away from, the target. We discarded trials in the incorrect direction (pro-reach: 2.8±2.1%; anti-reach: 5.9±3%) and with RT<140 ms (2.4±2.9%).

#### EMG data recording

Surface EMG activity was recorded from eight upper-limb muscles including four monoarticular shoulder muscles (clavicular head of the pectoralis major, PMch; sternal head of the pectoralis major, PMsh; anterior deltoid, AD; posterior deltoid, PD), two bi-articular arm muscles (biceps brachii, BIC; long head of the triceps brachii, TRIloh) and two monoarticular elbow muscles (brachioradialis, BR; lateral head of the triceps brachii, TRIlah). Surface electrodes with inbuilt pre-amplification and filtering (10x, 20-450Hz; Delsys Inc. Bagnoli-8 system, Boston, MA, USA) were used to record the EMG signals. The raw signal was further amplified by 100x in the “Delsys Bagnoli-8 Main Amplifier Unit” and sampled at 2 kHz using a 16-bit analog-digital converter (USB-6343-BNC DAQ device, National Instruments, Austin, TX). The quality of the EMG signal was checked online with an oscilloscope. The EMG data were then filtered with a 20–300 Hz bandwidth filter and full-wave rectified during offline analysis.

#### Identification of express visuomotor responses

For the first experiment, EVRs were identified trial-by-trial via a procedure that we recently developed and validated ^31^. Briefly, we first down sampled the EMG data to 1 kHz and computed the trace integral between 100 ms before and 300 ms after the target onset time. We then detrended the integrated signal by subtracting the linear regression function of the background period (i.e. from 100 ms before to 70 ms after the stimulus presentation) from the entire 400 ms window. We indexed the “candidate” muscle response onset time as the first time the detrended-integrated signal exceeded the average background value by more (i.e. earliest muscle activation), or less (i.e. earliest muscle inhibition), than five standard deviations. Finally, we ran a linear regression analysis around the candidate muscle response onset time and indexed the time point at which the linear trendline intercepted the zero value in the detrended-integrated signal. Consistent with previous work, we classified a muscle response as “express” if it was initiated within 70-120 ms after the target presentation ^24,25,31,40^. Results from experiment 1 relied on the subset of trials in which EVRs were detected via this method.

Note that the anti-reach task of the second experiment required directing the reach toward a non-veridical location opposite to the visual stimulus location. Consistent with previous work ^25^, we found smaller express responses in the anti-target than pro-target reaching conditions. Given this, only a limited proportion of express responses could be detected via the trial-by-trial detection method. Thus, to quantify any subtle changes in muscle activity from baseline within the express response time in the second experiment, we simply took the mean EMG relative to baseline for all trials within a pre-defined time period (90-100ms post stimulus). We used this narrow window, that coincided approximately with the mean EVR onset time in experiment 1, to create a conservative measure of EVR signals that was least likely to be contaminated by LLR activity. EVR results from experiment 2 therefore relied on the mean EMG measured in the EVR window for all trials.

#### Target-dependent muscle activity and preferred reaching direction

For the first experiment, the target-dependent express muscle activity was computed by normalizing the response magnitude across the eight target directions. More precisely, for each detected express-response trial, we computed the express response magnitude by subtracting the average background value from the average EMG activity recorded in the 10ms after the muscle response initiation time. We then normalized the target-dependent response amplitudes to the largest and smallest values across the eight target locations, such that the tuning curve ranged ±1 for each muscle and subject. On the same trials, we also computed the target-dependent muscle activity in the long-latency epoch by taking the EMG signal recorded during the 10ms before the mechanical RT. For both express and long-latency visuomotor responses, we also defined the muscle preferred reaching direction (PRD) by computing the vector sum of the eight target-oriented express and long-latency muscle responses.

For the second experiment, the express response magnitude was calculated across all correct trials (see previous section) by averaging the EMG signal enclosed between 90ms and 100ms after the target presentation, which was the time-window in which most express responses were most prominent in the first experiment. For each task condition, we then averaged the magnitude of express and long-latency muscle responses across all trials.

#### Muscle coordination analysis

This study tested the idea that a putative subcortical sensorimotor network can compute musculoskeletal dynamics strategies needed to drive goal-directed reaches. If so, then EVRs should reflect the same muscle synergies as those recruited to voluntarily reach the target (LLRs). To this aim, we calculated the muscle synergies using PCA. For the first experiment, we initially defined a low-dimensional subspace of muscle synergies from EVRs in the abducted-shoulder posture condition. Specifically, for each possible number of synergies (i.e. 1:7), we ran the PCA to decompose the reference target by muscle (8×8) matrix and then reconstruct the target-dependent muscle activity. Subsequently, we computed the “*variance accounted for”* (VAF) score between the original and reconstructed muscle activity and took the minimum number of components (4) that returned >95% VAF across all subjects. Finally, we used the low-dimensional subspace of EVR-derived muscle synergies to reconstruct the LLR muscle activity in the abducted posture condition, as well as the EVR and LLR muscle activity recorded in the adducted posture condition. We reasoned that if the express and long-latency responses reflect similar sensory-to-motor transformation of the hand-to-target position, then the low-dimensional subspace should be more accurate to reconstruct the muscle activity across responses than postures.

For the second experiment, we assessed how well EVR and LLR muscle activity observed in the pro-reach task could be reconstructed via a low-dimensional subspace of muscle synergies that was computed from EVRs for the same stimulus-locations during the anti-target reaches. We reasoned that if a subcortical sensorimotor network can coordinate muscles for a reaching action, then EVR muscle synergies generated by presentation of a visual stimulus in the anti-reach task should accurately reconstruct the muscle activity required to reach targets, even when the subsequent reach is directed in the opposite direction from the stimulus.

#### Statistical analysis

Standard statistical analyses were performed in Jamovi (v. 2.2.5). Results were analysed with repeated measures ANOVA analysis with Bonferroni correction (normality of the distributions was verified by the Shapiro–Wilk test), unless otherwise stated. Where appropriate, we ran Bonferroni corrected post-hoc tests. For all tests, the statistical significance was designated at P < 0.05. Circular statistics were calculated in Matlab using the CirsStat toolbox ^87^. For both the express and long-latency visuomotor responses, we defined the muscle preferred reaching direction by computing the vector sum of the eight target-dependent muscle responses. Circular statistics were run on each muscle sample and response epoch to test the influence of initial upper-limb posture on the muscle tuning direction. Specifically, we calculated the circular mean and 95% confidence interval of the muscle preferred direction, the circular variance as 1-R (where R = resultant vector length) such that circular variance values close to 0 indicate uniform preferred directions with respect to the circular mean. To test whether the muscles were directionally tuned, we ran both the Rayleigh and Omnibus uniformity tests that assess whether the muscle preferred reaching directions could have been drawn from a normal Von Mises distribution. Finally, we tested whether differences in muscle tuning direction between the two postures were statistically significant via the Watson-Williams test.

## Acknowledgements

This work was supported by an operating grant from the Australian Research Council (DP240101968) awarded to T.J. Carroll, S. Contemori, B.D. Corneil, G.E. Loeb and G. Wallis.

## Supplementary materials

**Supplementary figure 1.**
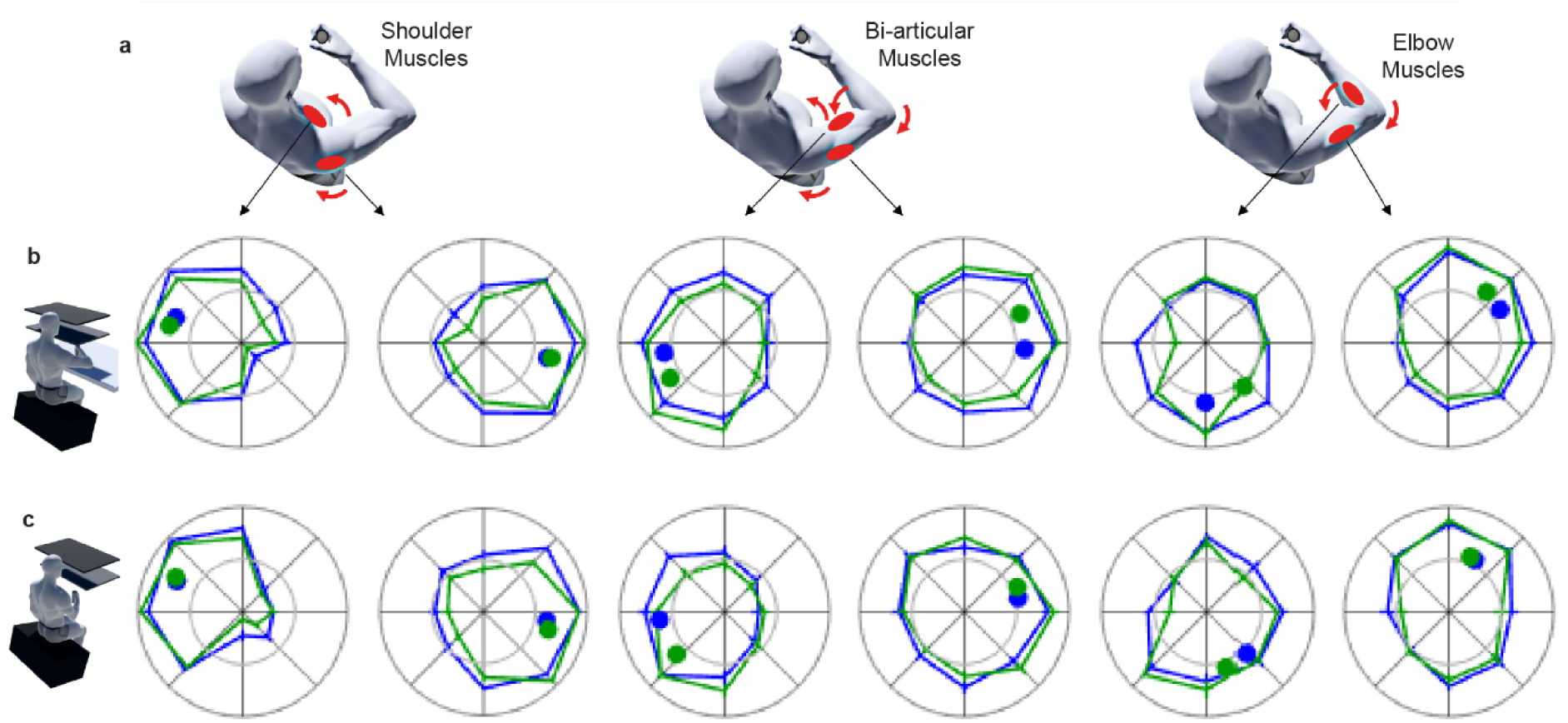
Responses when target muscle is unloaded. **a**, Schematic of the location and function of the six muscles used to illustrate target- and posture-specific muscle activation patterns for express and long latency muscle responses. **b**, Express (blue) and long-latency (green) muscle activity for each target direction in the abducted posture when muscles were not loaded prior to target presentation. Each radial plot shows average (+/- SE) target-dependent tuning curves across the twenty participants. Express and long-latency activation values are normalised to the maximal EVR and LLR amplitudes respectively across target directions. The outer ring indicates the normalised value (1) and the inner ring represents the reference background activity (0) for each direction. Thus, excitations with respect to baseline lie between the two rings, and inhibitions from baseline fall within the inner ring. Note that inhibitions are minimal due to a lack of pre-loading forces. Blue (EVR) and green (LLR) circles show the preferred direction of each tuning curve. **c**, Same as **b**, except for the adducted posture.

**Supplementary figure 2.**
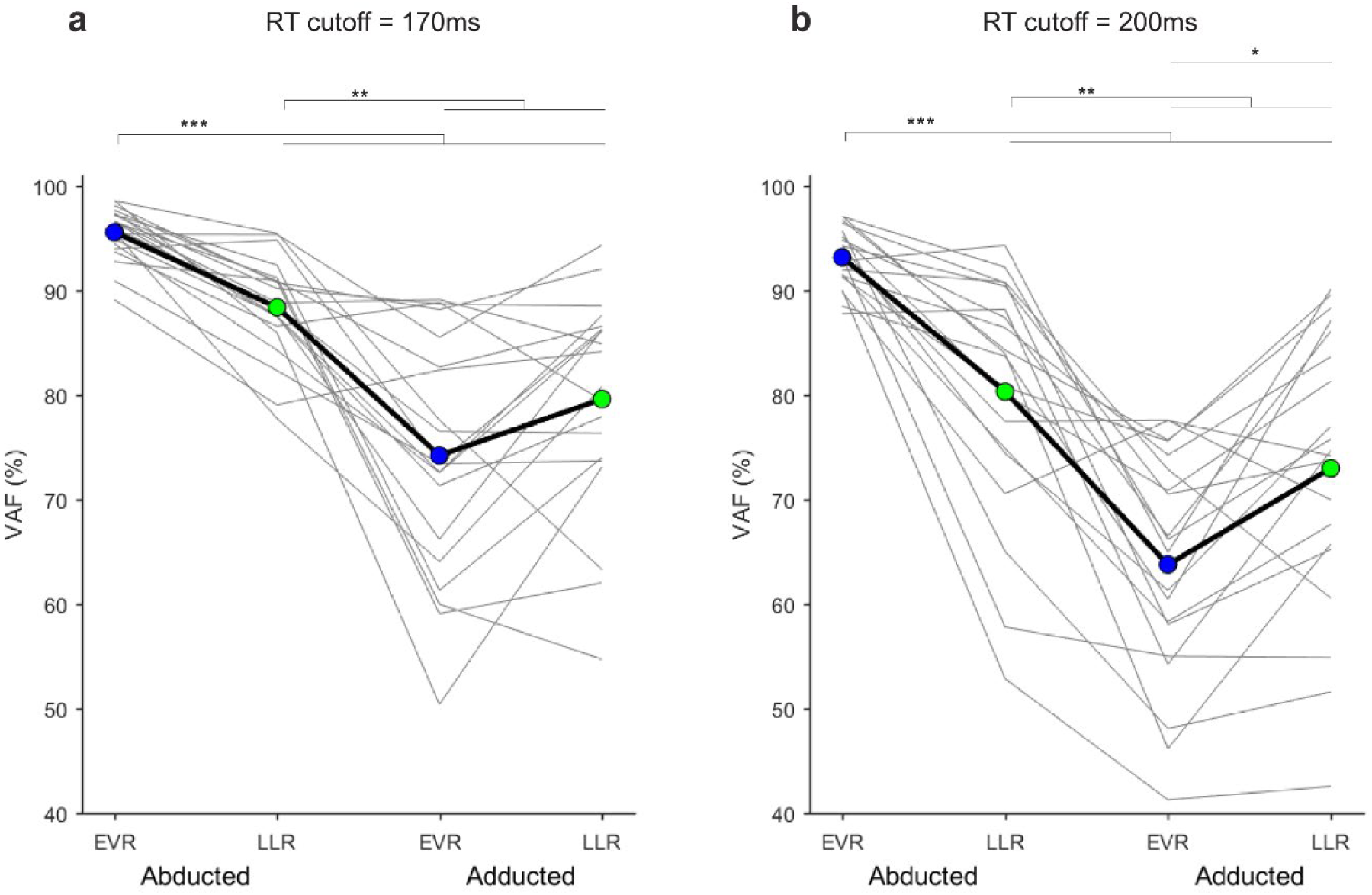
PCA results for alternative reaction time exclusion thresholds in Exp 1. PCA summary data for 170ms and 200ms reaction time cutoffs for EVR inclusion. **a,** Variance accounted for (VAF) by the top 4 principal components extracted from EVRs, extracted when trials with reaction times less than 170ms were excluded, across stimulus directions in the abducted shoulder posture, for LLRs in both postures and EVRs in the adducted posture in experiment 1. **b,** VAF by the top 4 principal components extracted from EVRs, extracted when trials with reaction times less than 200ms were excluded, across stimulus directions in the abducted shoulder posture, for LLRs in both postures and EVRs in the adducted posture in experiment 1. EVR means are blue, LLR means are green. The brackets and asterisks indicate the statistically significant (* *p*<0.05, ** *p*<0.01, *** *p*<0.001,) post-hoc pairwise contrasts.

**Supplementary figure 3.**
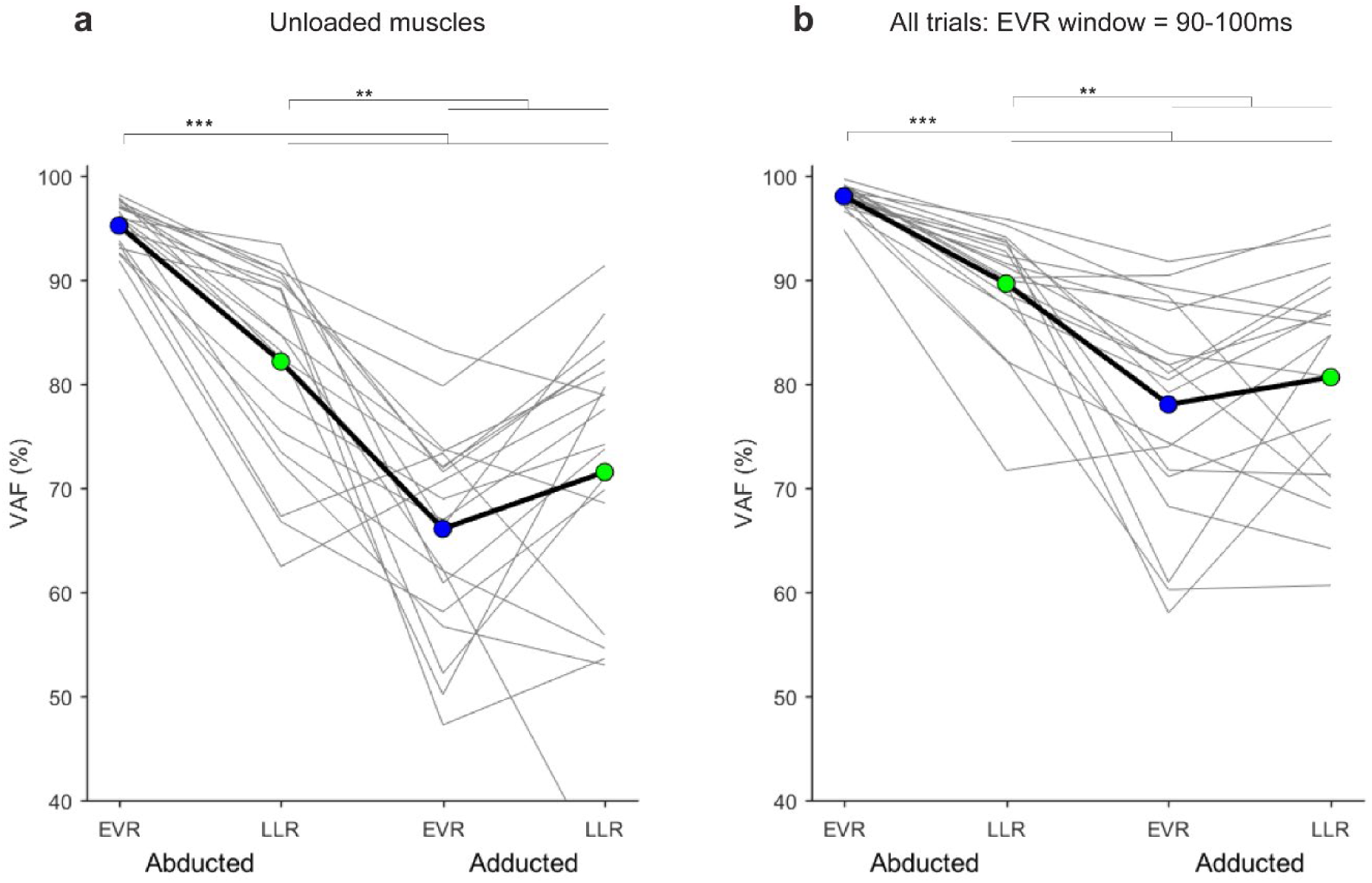
PCA results for alternative EVR measures in Exp 1. **a,** PCA summary data for EVRs extracted from muscles that were not pre-loaded. Variance accounted for (VAF) by the top 4 principal components extracted from EVRs, across stimulus directions in the abducted shoulder posture, for LLRs in both postures and EVRs in the adducted posture in experiment 1. **b,** PCA summary data for average EMG recorded in the EVR window for all trials, irrespective of whether an EVR was identified by our onset detection algorithm. VAF by the top 4 principal components extracted from EVRs in the abducted shoulder posture, for LLRs in both postures and EVRs in the adducted posture in experiment 1. EVR means are blue, LLR means are green. The brackets and asterisks indicate the statistically significant (* *p*<0.05, ** *p*<0.01, *** *p*<0.001,) post-hoc pairwise contrasts.

**Supplementary figure 4.**
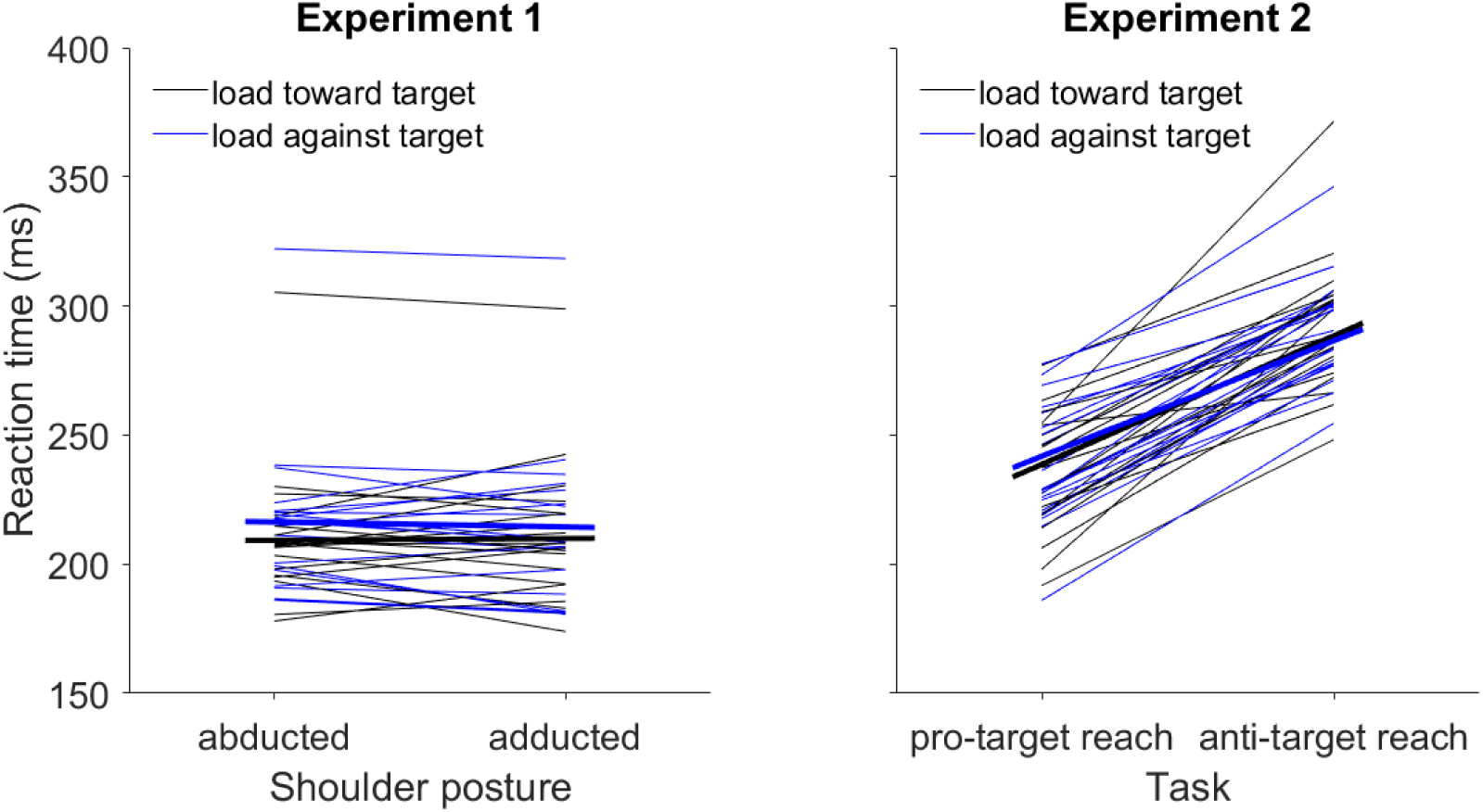
Mean reaction times by condition. Dependency of the volitional RT on (A) the postural conditions of experiment 1 and (B) the task conditions of experiment 2. For each plot, the thin lines represent single participants, and the thick lines represent the average across participants. The black and blue lines represent the conditions in which the robot force was directed toward and against the target, respectively.

## REACTION TIME ANALYSIS

**Experiment 1**: RM 2-way ANOVA with posture (abd and add) and preload direction (toward target and against target) as within subject factors. The RT was significantly modulated by the preload direction (F_1,19_=20.74, *p* <0.001) and posture*preload interaction (F_1,19_=4.66, *p* = 0.044). The post-hoc analysis showed that the RT was significantly earlier when the robot force was directed toward the target rather than away from the target for both the abducted (*p*<0.001) and adducted (*p*=0.033) shoulder postures.

**Experiment 2**: RM 2-way ANOVA with task (pro-target and anti-target reaches) and preload direction (toward target and against target) as within subject factors. The RT was significantly modulated by the task condition (F_1,19_=224.45, *p* <0.001). The post-hoc analysis showed that the RT was significantly later (*p*<0.001) when the reach was directed away from the target rather than toward the target.

---

**Supplementary table 1.** EVR prevalence and onset times. Muscle-dependent prevalence and onset time of express visuomotor responses for experiment 1. The data are shown as mean (standard deviation). Abbreviations: PEC-ch, pectoralis major clavicular head; PEC-sh, pectoralis major sternal head; AD, anterior deltoid; PD, posterior deltoid; BIC, biceps brachii; TRI-loh, triceps brachii long head; BR, brachioradialis; TRI-lah, triceps brachii lateral head.

| Muscle | EVR prevalence (%) | EVR onset time (ms) |
| --- | --- | --- |
| PEC-ch | 59 (7.5) | 89 (3.4) |
| PEC-sh | 55 (10) | 88 (2.7) |
| AD | 51 (7.1) | 92 (2.4) |
| PD | 56 (6.7) | 92 (2.9) |
| BIC | 46 (5.9) | 93 (2.8) |
| TRI-loh | 48 (5.7) | 91 (2.6) |
| BR | 46 (5.7) | 92 (2.5) |
| TRI-lah | 49 (5.6) | 92 (3.1) |

**Supplementary table 2.**
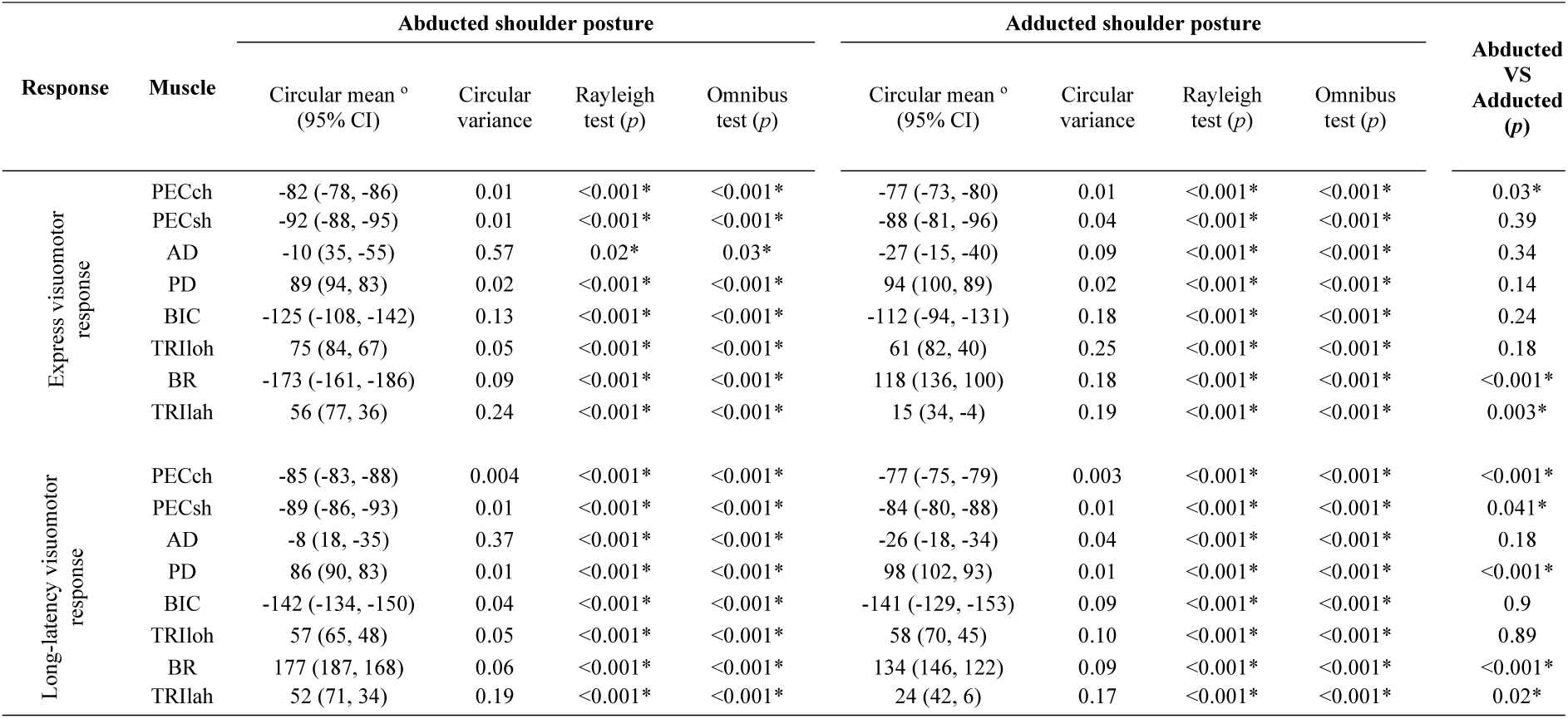
Circular statistics summary for muscle preferred directions. Dependency of the muscle tuning direction on the response epoch and initial posture condition. The asterisks indicate statistically significant findings. Abbreviations: CI, confidence interval; PECch, pectoralis major clavicular head; PECsh, pectoralis major sternal head; AD, anterior deltoid; PD, posterior deltoid; BIC, biceps brachii; TRIloh, triceps brachii long head; BR, brachioradialis; TRIlah, triceps brachii lateral head.

## Results

Table S2 shows the summary results of the circular statistics analysis. For all muscles, response epochs, and postural conditions we found a circular variance <0.5, except for the anterior deltoid EVR in the abducted shoulder posture (0.57). These results indicate that the muscle activity was consistently tuned across the eight different target directions and were corroborated by the statistically significant results of the Rayleigh and Omnibus test, thus showing that the muscles preferred direction was clustered around the circular average.

We found that the directional tuning of five muscles changed across the posture condition, and the change was in the same direction for both the express and long-latency responses (Table S2). For three of these muscles, the effect of posture on the tuning direction was statistically different for both the express and long-latency epochs; for two other muscles, a significant effect of posture was observed only for the long-latency epoch.

## Notes

### Competing Interest Statement

The authors have declared no competing interest.

