## Supplementary figures and images for "Subcortical control of reaching in humans"

### Supplementary Figure 1

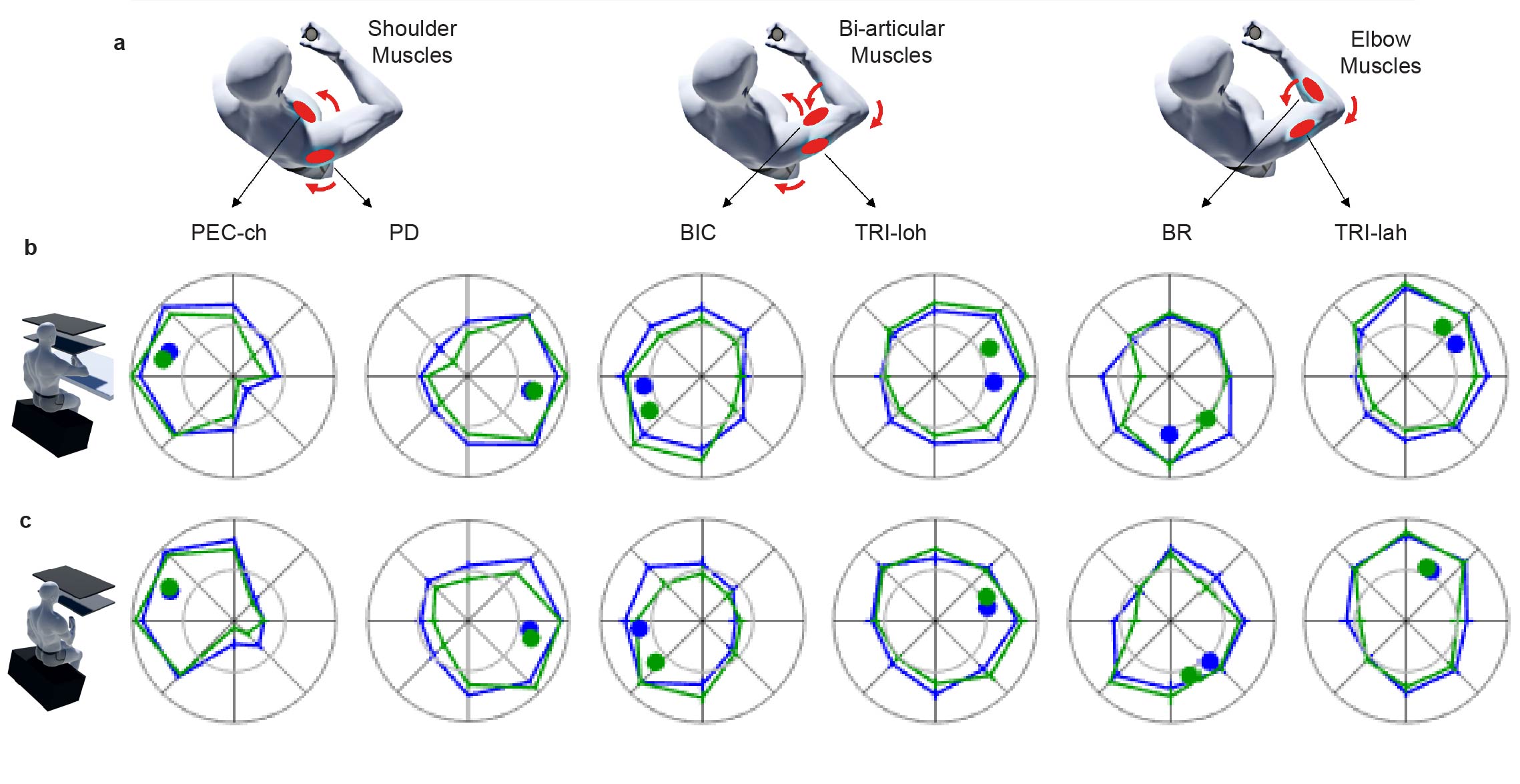

### Supplementary Figure 2

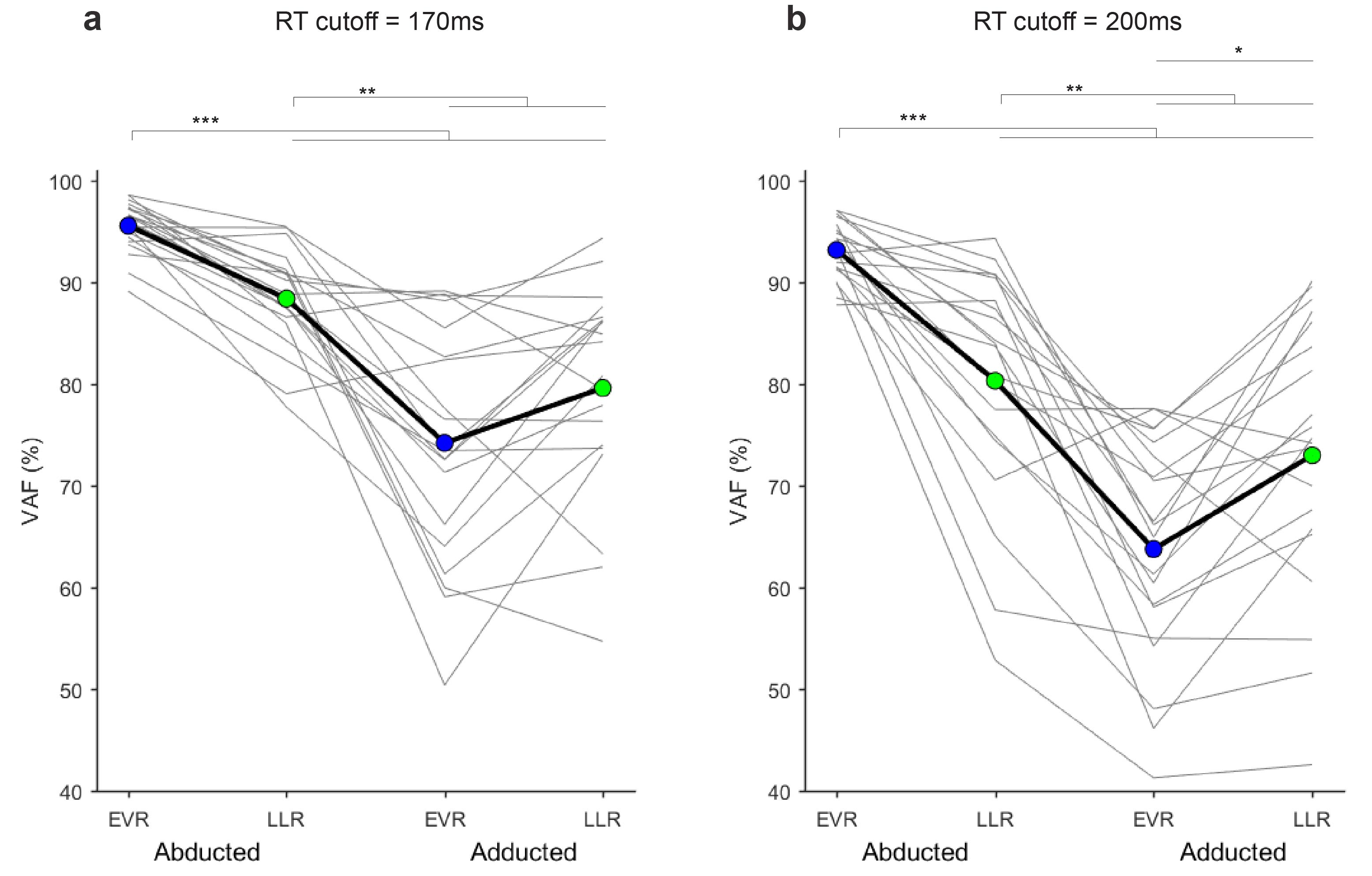

### Supplementary Figure 3

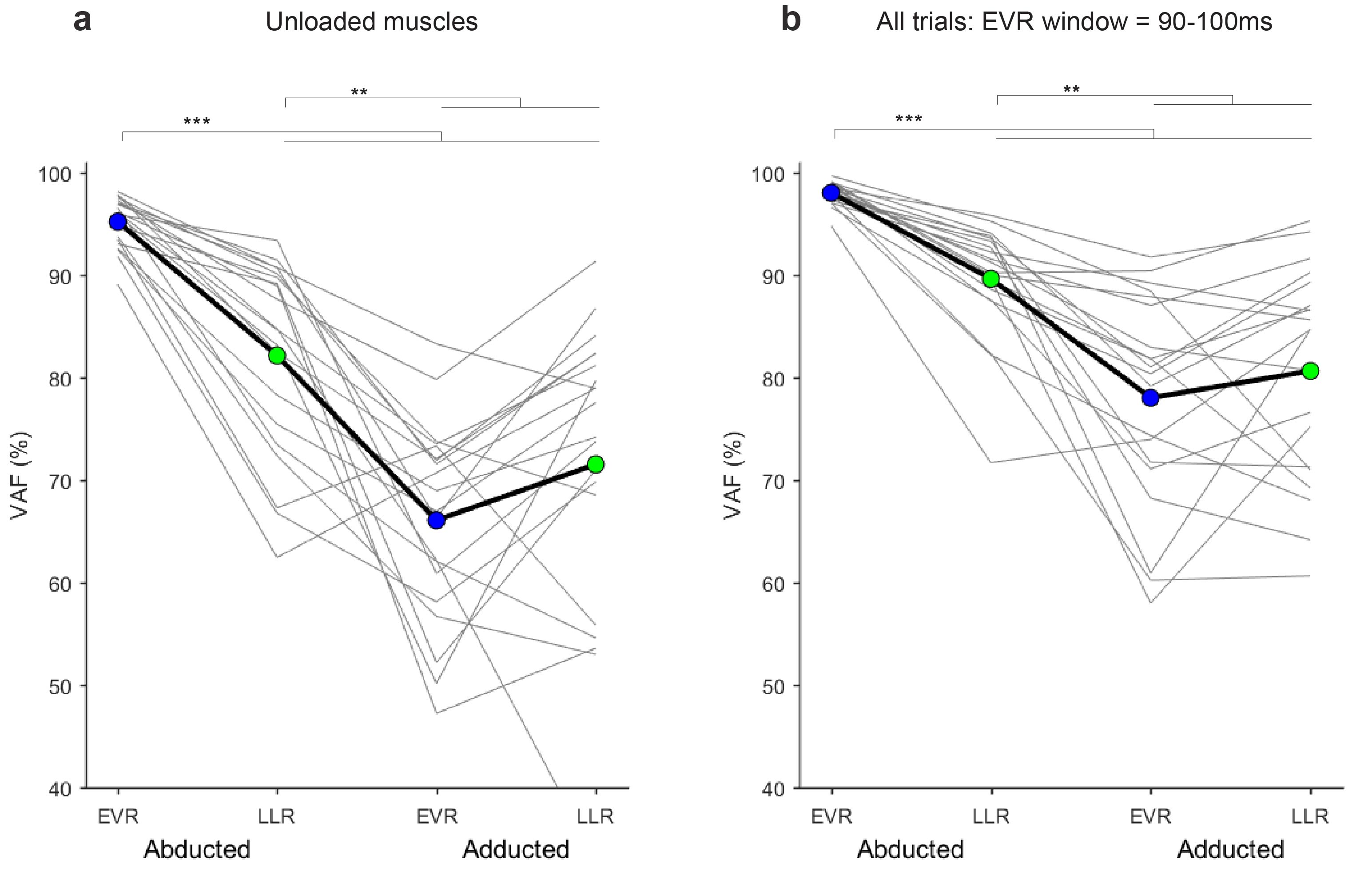

### Supplementary Figure 4

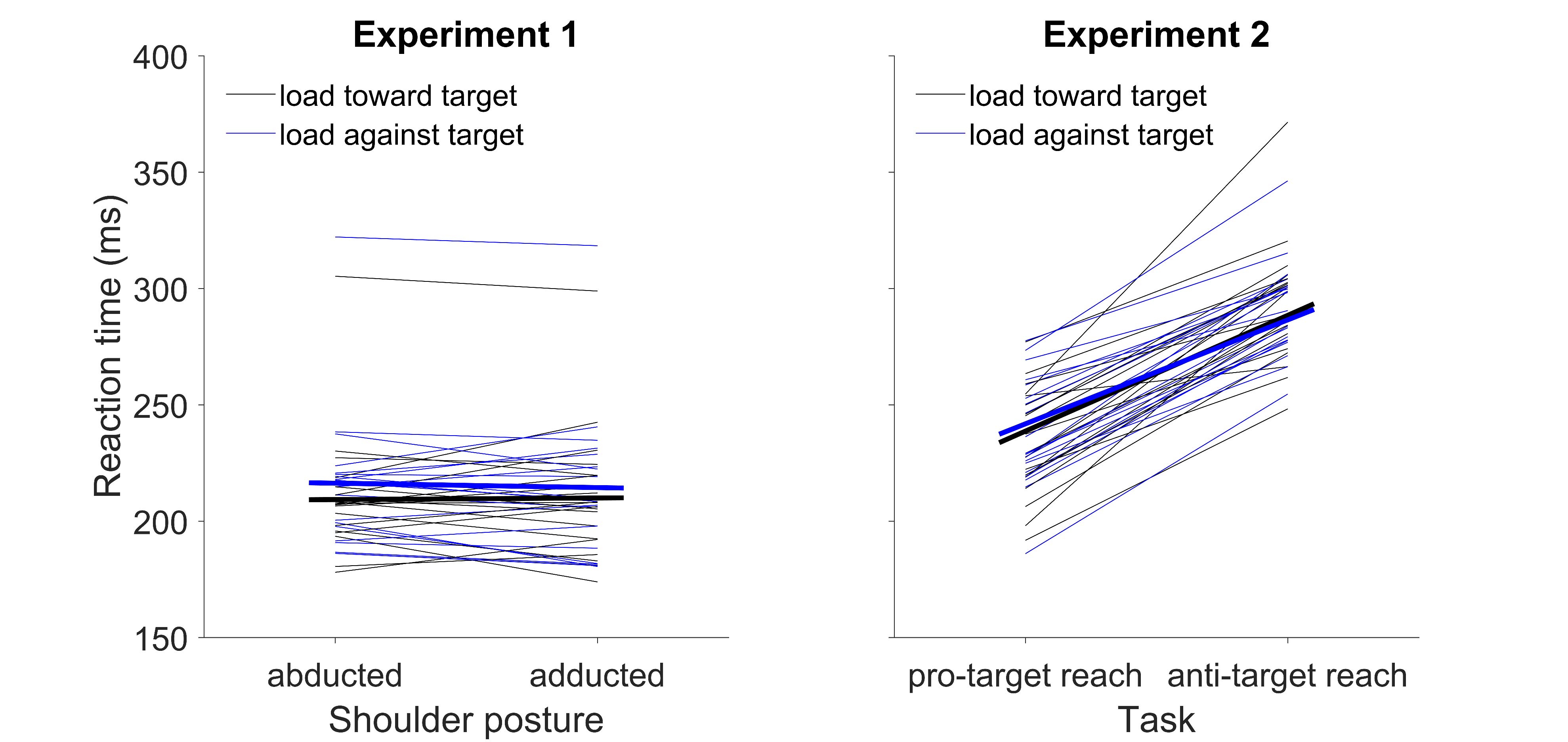
